# Genetic atlas of hygro- and thermosensory cells in the vinegar fly *Drosophila* melanogaster

**DOI:** 10.1101/2023.06.26.546511

**Authors:** Kristina Corthals, Vilma Andersson, Allison Churcher, Johan Reimegård, Anders Enjin

## Abstract

The ability of animals to perceive and respond to sensory information is essential for their survival in diverse environments. While much progress has been made in understanding various sensory modalities, the sense of hygrosensation, which involves the detection and response to humidity, remains poorly understood. In this study, we focused on the hygrosensory, and closely related thermosensory, systems in the vinegar fly *Drosophila melanogaster* to unravel the molecular profile of the cells of these senses. Using a transcriptomic analysis of over 37,000 nuclei, we identified twelve distinct clusters of cells corresponding to temperature-sensing arista neurons, humidity-sensing sacculus neurons, and support cells relating to these neurons. By examining the expression of known and novel marker genes, we validated the identity of these clusters and characterized their gene expression profiles. We found that each cell type could be characterized by a unique expression profile of ion channels, GPCR signaling molecules, synaptic vesicle cycle proteins, and cell adhesion molecules. Our findings provide valuable insights into the molecular basis of hygro- and thermosensation. Understanding the mechanisms underlying hygro- and thermosensation may shed light on the broader understanding of sensory systems and their adaptation to different environmental conditions in animals.

## Introduction

The astonishing diversity of sensory systems among animals enables them to navigate and survive in an array of challenging environments. In recent decades, significant progress has been made in unraveling the underlying mechanisms of many of these sensory modalities, however, a few senses remain unknown. One of them is hygrosensation – the ability to detect and respond to humidity. Humidity is a climactic factor crucial for the survival of terrestrial animals [1,2]. Insects in particular are sensitive to variations in humidity and use this ability to guide them in specific behaviors such as food-seeking and oviposition site-selection [3,4]. Humidity is detected by humidity receptor neurons (HRNs), which are located within a specialized sensory structure called the hygrosensillum [5]. In insects, hygrosensilla are found on the antenna, a pair of appendages on the head that functions as a multi-sensory organ detecting temperature, odorants and sound in addition to humidity. The hygrosensilla are intermingled with olfactory sensilla and shares their general structure. However, unlike olfactory sensilla, which have pores to enable odorant molecules to reach the sensory receptors, hygrosensilla are covered by a poreless hydrophobic cuticle. As a result, the HRNs are protected from exposure to airborne molecules thereby conferring an enigmatic attribute to the mechanism by which humidity is sensed. Typically, hygrosensilla consist of a moist neuron (sensitive to increased humidity), a dry neuron (sensitive to decreased humidity), and a hygrocool neuron (sensitive to decreased temperature), which are co-located within a sensillum, forming a hygrosensory “triad.”

In the vinegar fly *Drosophila melanogaster*, the hygrosensilla are found within an invagination of the posterior side of the antenna called the sacculus (Figure 1A-B) [6–9]. The sacculus is composed of three chambers, with the initial two dedicated to the detection of humidity while the third chamber contain olfactory receptor neurons (ORNs) that respond to acid and ammonium [10,11]. HRNs express members of the ionotropic receptor (IR) family, dry neurons express Ir40a, while moist neurons express Ir68a. Hygrocool cells found within the first chamber of the sacculus express IR21a, while the hygrocool cells in the second chamber express Ir40a [6–9, 12]. Additionally, Ir21a is also expressed within the cold-sensing temperature receptor neurons (TRNs) found in the arista [13]. Hot cells in the arista express gustatory receptor GR28b.d [14]. HRNs and TRNs also express two additional IRs, Ir93a and Ir25a, which may act as co-receptors.

**Figure 1:**
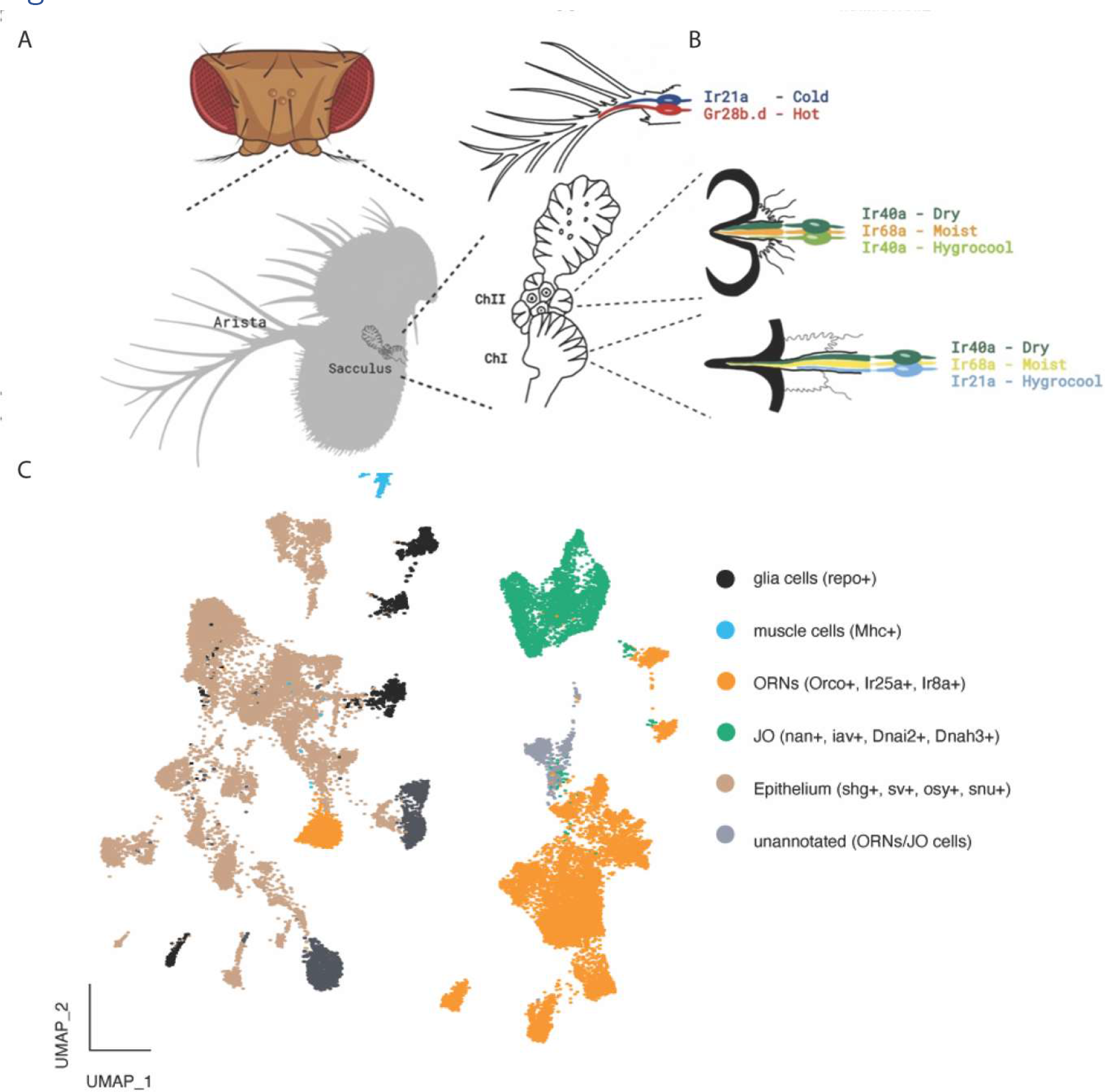
#A) Overview of Drosophila antenna. The sacculus is an invagination on the dorsal side of the antennal 3^rd^ segment. It consists of three chambers. Previous studies have located the hygrosensilla in chamber I and chamber II. The arista is a feather-like appendage on the antenna. Previous studies have located temperature sensitive cells in the arista. B) Sensory neurons the sacculus and the arista. Hygrosensory receptor neurons are found in two chambers of the sacculus, in so called hygrosensilla. Anatomical differences for the hygrosensilla in each chamber have been described previously. Each sensilla houses a triad of neurons, one moist cell, one dry cell and one hygrocool cell. In chamber I moist cells express Ir68a, dry cells Ir40a and hygrocool cells Ir21a. In chamber II moist cells express Ir68a, dry cells Ir40a and hygrocool cells Ir40a. The arista temperature cells are divided into hot cells (expressing Gr28b) and cold cells (expressing Ir21a). C) UMAP plot of the whole antennal data set (all segments + arista) with annotations for the different functional groups. ORN: olfactory receptor neurons. JO: Johnstons Organ. One cluster could not be assigned a definite identity, based on gene expression it is suspected to contain ORN and JO neurons.

Although all HRNs and TRNs express a sensory receptor protein, not all cells can be defined by the expression of a unique receptors suggesting other molecules are involved in the transduction mechanism. Here we describe an analysis of the antennal dataset from the fly cell atlas [15], consisting of > 37,000 nuclei. This analysis identified 1411 nuclei which we could identify as arista neurons, sacculus neurons or support cells in these structures. These clustered into twelve clusters which could be matched to both functional type and anatomical location. Our data show that dry, moist and hygrocool neurons in chambers I and II are transcriptomically distinct suggesting a functional distinction between these otherwise similar cell types. Furthermore, we identify several candidate genes that may have a role in the function of these cells, including the unknown mechanism of hygrosensory transduction.

## Results

### IDENTIFICATION OF ARISTA and SACCULUS CELLS

To build a transcriptomic atlas of the cells in the sacculus and arista we utilized the publicly available Fly Cell Atlas [15]. We started with the available raw data and performed a principal component analysis (PCA) followed by a Nearest Neighbor Clustering using the Seurat pipeline [16,17]. Clustering of all cells resulted in 36 Clusters (Supplemental Figure 1). Based on differential gene expression in these clusters an we can divide them into 5 functional groups (Figure 1C): glia cells, marked by the expression of *repo* [18], muscle cells, marked by expression of *Mhc* [19], olfactory/temperature/humidity receptor neurons marked by expression of co-receptors *Orco*, *Ir25a* and *Ir8a* [20,21]; Johnston’s organ (JO) cells marked by expression of *nan*, *iav*, *Dnai2* and *Dnah3* [22], epithelium and support cells marked by expression of *shg, sv*, *snu* and *osy* [23–25]. Similar to the Fly Cell Atlas annotation we are also left with one unidentified cluster. Differential gene expression of highly expressed genes of the unknown cluster shows close similarity to the expression pattern in the clusters assigned as JO cells.

To locate the cells of the sacculus and arista in the dataset we made use of the known expression of Ir93a, Ir40a and Ir21a in these cells [6–9]. We find combined expression of these receptors in only one cluster of the whole antennal dataset, cluster 15 (Figure 2A). The same cluster also shows expression of arista hot cell marker Gr28b and Ir64a, an acid sensor found in ORNs in sacculus chamber III (Figure 2A) [10,26]. In chamber III of the sacculus, another olfactory neuron sensing ammonium is also found, defined by the expression of RH50 and Amt [11]. These sacculus cells were found in another cluster, cluster 35.

**Figure 2:**
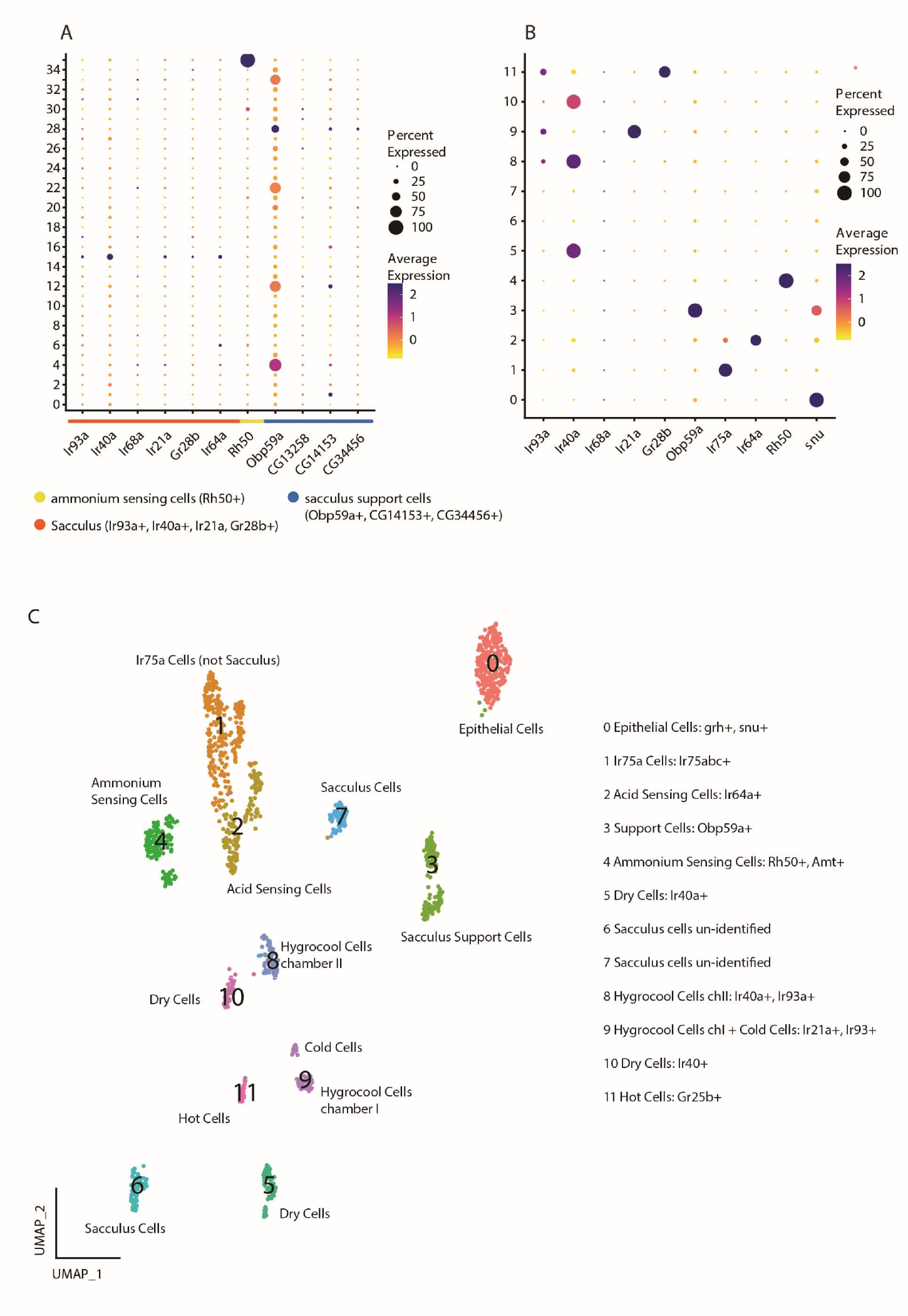
#A) Selection of clusters from full antenna dataset for sub-clustering. Dot plot shows expression of known marker genes: expression of Ir93a, Ir40a, Ir21a, Gr28b and Ir64a for sacculus (chamber I, II and III) and arista cells. Rh50 for ammonium sensing cells (sacculus III). Obp59a, CG13258, CG14153 and CG34456 for sacculus support cells. Based on these expression patterns cluster 15, 28 and 35 were selected for sub-clustering. B) Dot plot of known marker genes in the sub-clustering resulting in 12 clusters. All genes have previously been reported as marker genes for cell identities. C) UMAP projection of the 12 clusters with assumed identities based on gene expression from B.

Apart from neurons, the sacculus and arista sensilla also contain support cells. To identify the sacculus support cells we searched for the known markers Obp59a, CG13258, CG14153 and CG34456 [27,28]. While several clusters express Obp59a, at least at low levels, only cluster 28 shows combined expression of Obp59a, CG14153 and CG34456 (Figure 1B).

We extracted the cells from clusters 15, 28 and 35 and sub-clustered them using the Seurat pipeline. This resulted in 12 clusters (Figure 2 B). To define these clusters, we looked for expression of previously described markers and could identify the following (Figure 2 C): cluster 0, epithelial cells (based on expression of snu and grh [24,29]); cluster 1, acid sensing cells outside the sacculus (based on expression of Ir75a [21]); cluster 2, acid sensing cells in sacculus chamber III (based on expression of Ir64a [10]; cluster 3, sacculus support cells (based on expression of Obp59a [27,28]); cluster 4, ammonium sensing cells in sacculus chamber III (based on expression of Rh50 and Amt [11]); clusters 5, 8 and 10, dry cells in sacculus chamber I, II and hygrocool cells in chamber II (based on expression of Ir40a [6–9]); clusters 6 and 7: unknown; cluster 9: cold cells in arista and hygrocool cells in sacculus chamber I (based on expression of Ir93a and Ir21a [6–9]); cluster 11, hot cells of the arista (based on expression of Gr28b [14]). Clusters 6 and 7 likely correspond to moist cells in chamber I and II, however, Ir68a, a receptor previously described as expressed in moist cells [6–9], cannot be found in this sub-clustering and only in 18 cells in the whole antennal dataset. We are therefore disregarding it as a marker gene in this dataset (see Supplemental Figure 2).

### Validation of arista and sacculus cells

To describe the cells of the arista and sacculus in more detail, we extracted the top markers for each cluster and analyzed the relationship between the clusters based on the expression of these genes. This analysis showed that the clusters have pairwise relations: temperature-sensing cells in cluster 9 and 11, putative moist cells in cluster 6 and 7, IR40a-cells in clusters 5 and 10 (and to a smaller degree the IR40a-cells in cluster 8), olfactory cells in clusters 1, 2 and 4, and support cells and epithelial cells (Figure 3A).

**Fig 3:**
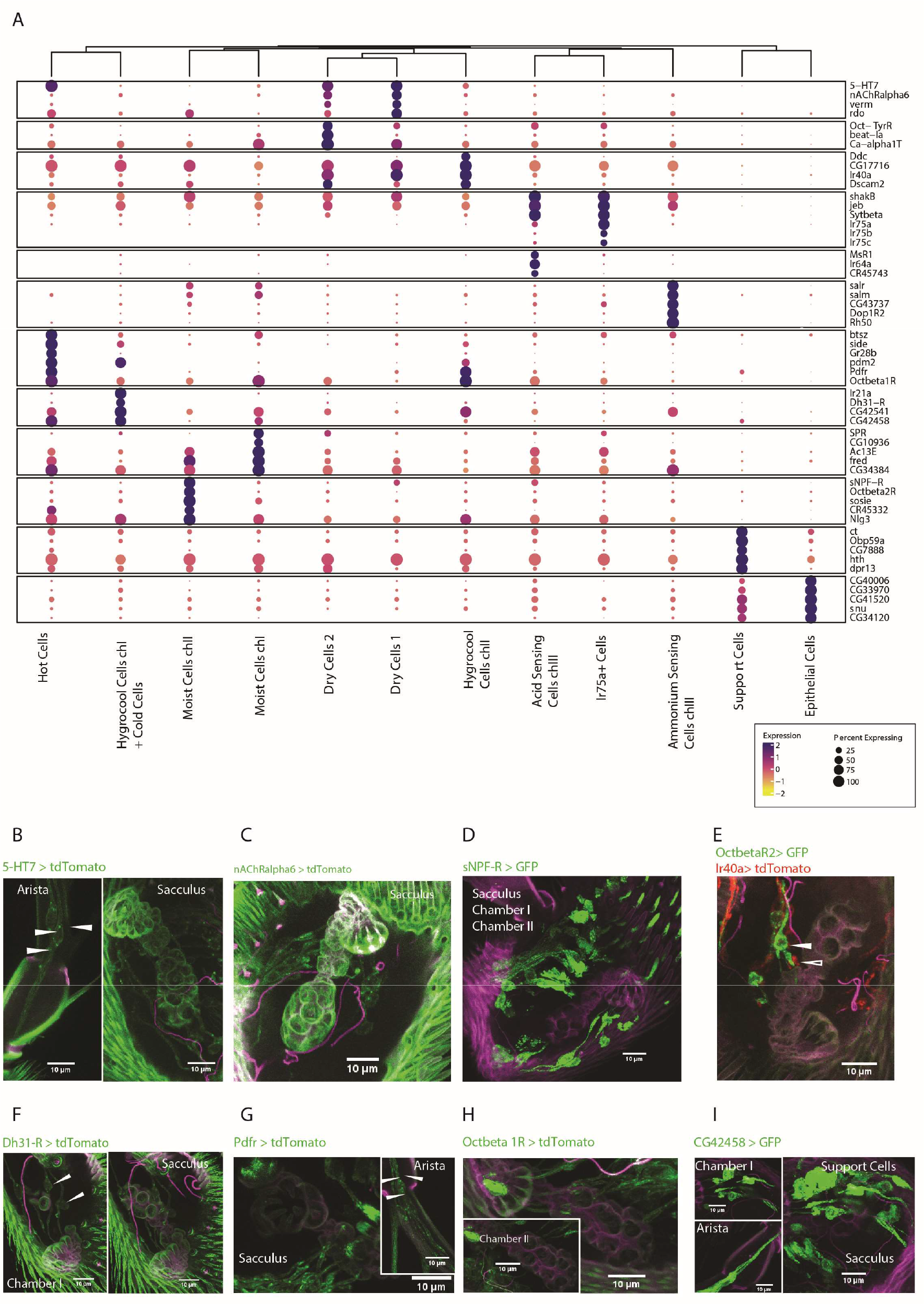
#A) Clustered dot plot showing the expression of top marker genes for each cluster. The dendrogram indicates the relationship between clusters based on this expression. B-I) Immunostainings for selected genes. Gal4 lines for these genes were crossed to either GFP or tdTomato reporter lines (see Material&Methods for more information on fly strains).

Next, we looked at the expression of some of the novel markers identified to validate the identity of the clusters (Figure 3B). We used knock-in Gal4-lines generated to maintain endogenous regulatory sequences reporting the expression of selected genes crossed to tdTomato-reporters to determine the expression these genes. 5-HT7 and nAChRα6 are both found in cluster 5 and 10 and both genes were expressed in one cell per sensillum in chamber I and II. Additionally, GMR21D03-Gal4, a Gal4-line with expression driven from an enhancer of nAChRα6, showed an identical expression pattern in the antenna and stained the VP4 glomerulus in the brain which led us to conclude that clusters 5 and 10 contain dry cells in chamber I and II [30]. Since clusters 5 and 10 are composed of dry cells, cluster 8, the other cluster with expression of IR40a, contain hygrocool cells in chamber II. Pdfr also showed expression in cluster 8, and the hot cell cluster 11. Pdfr-Gal4 crossed to UAS-tdTomato stained cells in the arista and chamber II confirming the identity of cluster 8 as belonging to cells in chamber II. Cluster 6 shows specific expression of Octβ2R and staining for Octβ2R>tdTomato specifically labeled cells in chamber II (Figure 3). Co-staining with Ir40a showed that Octβ2R and Ir40a are not expressed in the same cells and cluster 6 therefore represents moist cells in chamber II. CG32458 is found in cluster 7, 9 and 11. The gene is expressed in the arista, corresponding to cluster 11, and in two cells per sensillum in chamber I. Since cluster 9 contain hygrocool cells in chamber I, cluster 7 likely corresponds to moist cells in chamber I. CG32458, and Pdfr, are both also expressed in support cells surrounding the sacculus corresponding to their expression in cluster 3.

Cluster 9 is a mix of cold cells of the arista and hygrocool cells. The top marker for this cluster, Ir21a, is expressed in both cell types. Expression analysis of another marker for this population, Dh-31R, showed expression in only chamber I, confirming that cluster 9 is a mixed population. To test if we could separate the cold cell and hygrocool cell in chamber I, we extracted the clusters with temperature-sensitive cells, clusters 8, 9 and 11 and performed a sub-clustering. This resulted in four clusters defined by the expression of Ir21a, Gr28b, Ir40a and Dh31-R respectively, suggesting they represent cold cells, hot cells and hygrocool cells in chamber II and I (Supp Figure 3).

### Genetic profile of arista and sacculus cells

Next, we looked at the genes defining each cluster (Supp Figure 4 – Supp Figure 8). To determine gene expression patterns that may underlie the functional differences between the cells in the arista and sacculus, we examined the expression patterns of four categories of genes important in shaping a neuronal phenotype: ion channels, GPCR signaling, synaptic vesicle cycle proteins and cell adhesion molecules. We used the top 100 genes defining each cluster using the average log2FC value and classified them with gene ontology, GLAD and Flybase terms using the PANGEA tool [31]. After correcting for genes present in more than one cluster, 819 genes were used in the analysis. Fourty-eight of these were classified as ion channels, 52 as involved in GPCR signaling, 68 involved in synaptic vesicle cycle and 34 as cell adhesion molecules.

Ligand-gated and voltage gated ion channels comprised the majority of ion channels. Ligand-gated channels showed the greatest specificity within the HRNs and TRNs (Figure 4). The glutamate receptors Glucalpha and GlurIB are enriched in dry cells and cold cells respectively while the nicotinic receptors nachalpha6 and nachalpha7 are enriched in dry cells and moist cells. The voltage-gated channels are more generally expressed between the neuronal clusters except for the Ca^2+^-channel Ca-alpha1T that is enriched in dry cells and K^+^-channel Shawl that is enriched in ammonium-sensing cells in chamber III. One TRP channels is enriched, Trpm, expressed by HRNs in chamber II.

**Fig 4:**
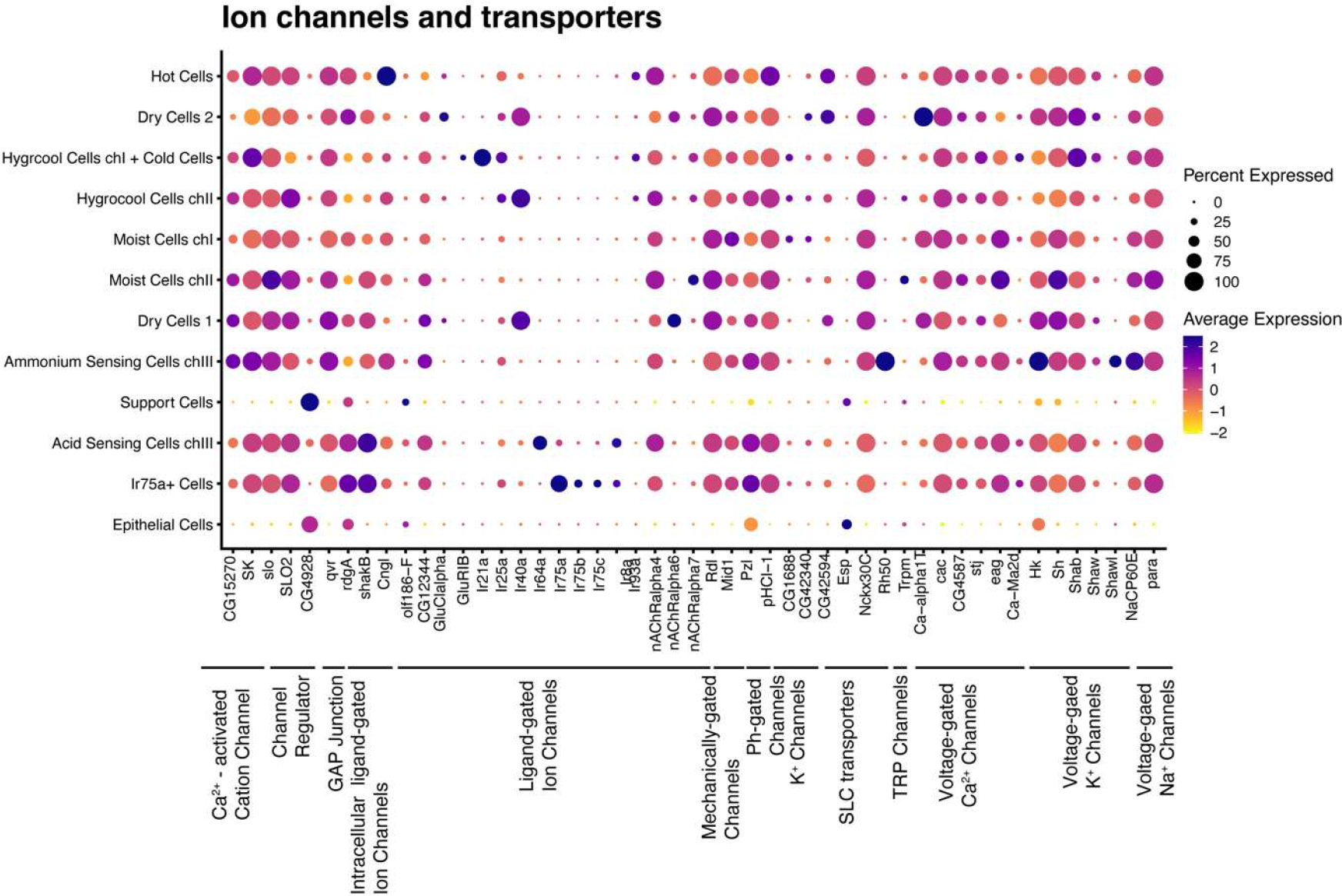
Ion Channels. Dot plot showing expression of selected ion channels. Classification of genes is indicated by the black lines. The size of the dot indicates the fraction of cells in each cluster expressing the genes. The average gene expression for each cluster is shown as a heatmap.

Of the GPCRs enriched, a majority are receptors for neurotransmitters (n=15) or neuropeptides (n=14) (Figure 5). The neurotransmitters include receptors for dopamine, GABA, acetylcholine, serotonin, tyramine and octopamine. Notably, all octopamine receptors annotated in the *D. melanogaster* genome are expressed in the sacculus in a subtype-specific pattern. Neuropeptide receptors are subtype-specific and most neuronal cell types in the arista and sacculus at have a unique expression of one or more neuropeptide receptors. In dry cells an orphan GPCR, CG43790, is specifically enriched. We also find specific expression of several intracellular components of G-protein signaling such as the adenylate cyclases Ac13E and Ac76E.

**Fig 5:**
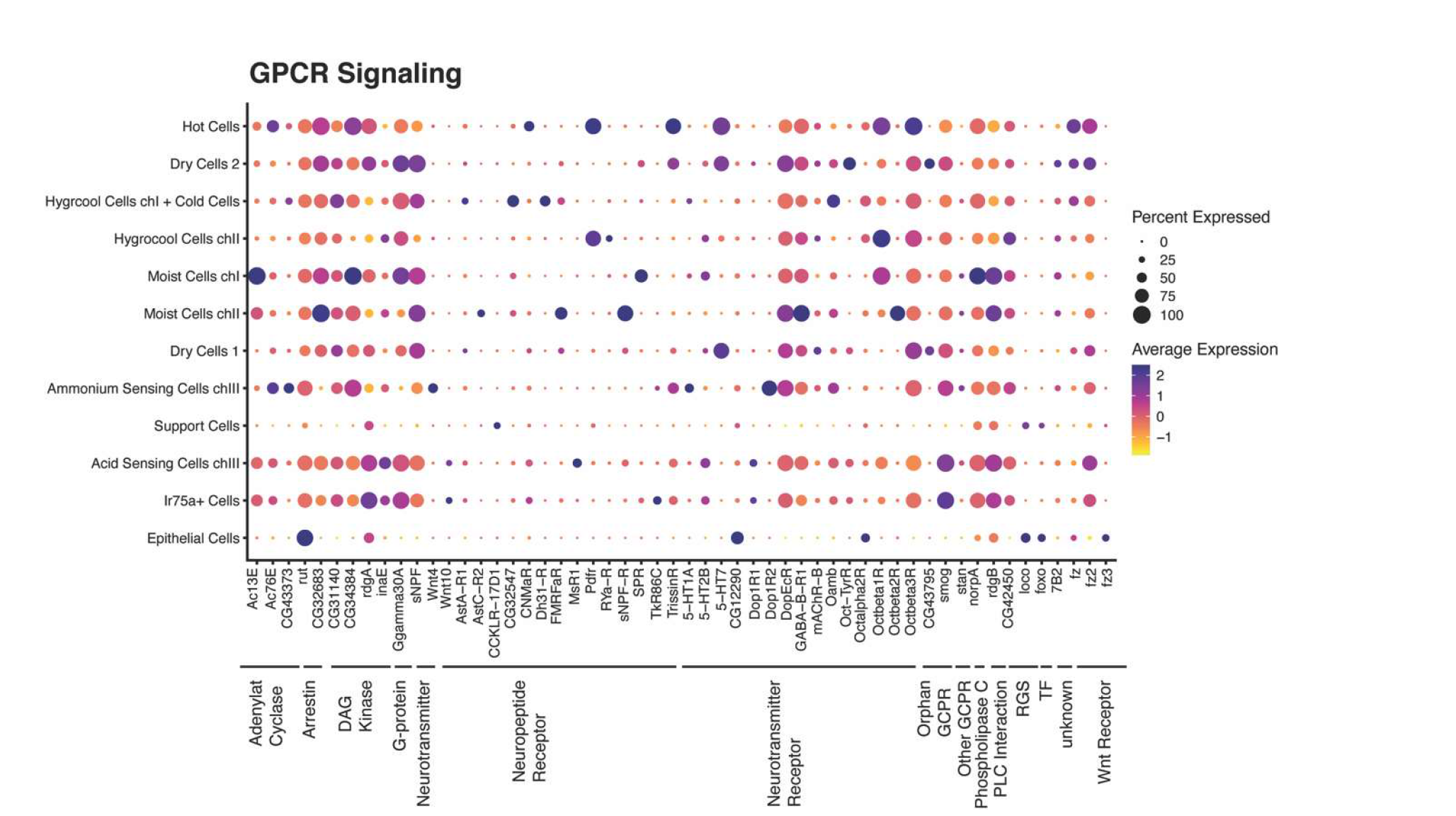
GPCR signalling. Dot plot showing expression of selected genes involved in G protein coupled receptor signalling. Classification of genes is indicated by the black lines. The size of the dot indicates the fraction of cells in each cluster expressing the genes. The average gene expression for each cluster is shown as a heatmap.

Genes relating to the synaptic vesicle machinery make up the largest number of genes among those defining the separate clusters (Figure 6). Fourteen of these genes belong a family of synaptic adhesion molecules called Defective proboscis extension response (Dpr) and Dpr-interacting proteins (DIPs). Different members of Dprs and DIPs form hetero- and homophilic interactions with each other and are expressed in a subtype-specific pattern. Another family of synaptic adhesion proteins that show a subtype-specific pattern are the neuroligins with a specific enrichment of Nlg2 in dry cells and Nlg4 in temperature-sensing cells. We also see an enrichment of the neuronal calcium sensor Frq2 in moist cells and hygrocool cells in chamber II.

**Fig 6:**
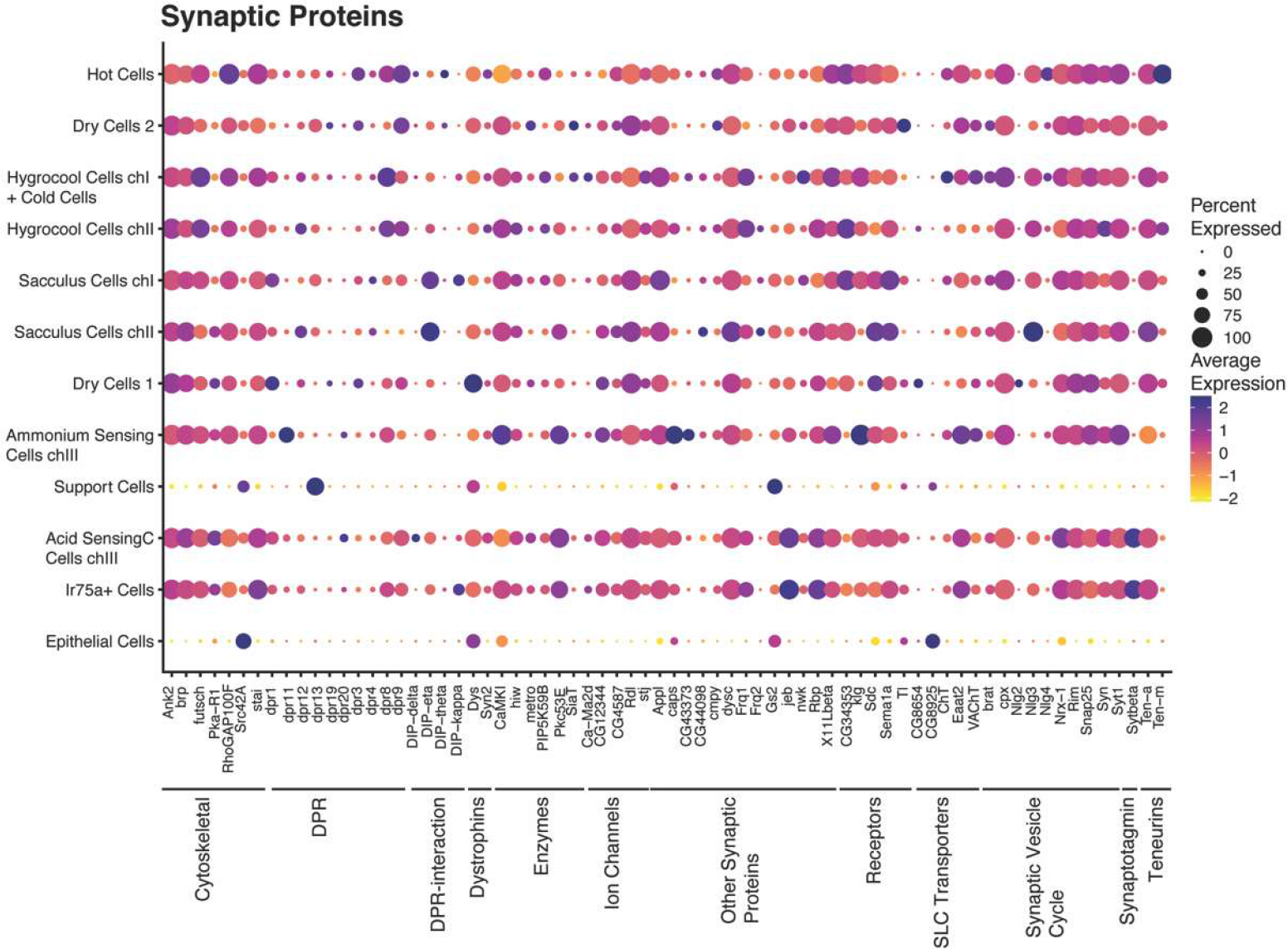
Synaptic Proteins. Dot plot showing expression of selected synaptic proteins. Classification of genes is indicated by the black lines. The size of the dot indicates the fraction of cells in each cluster expressing the genes. The average gene expression for each cluster is shown as a heatmap.

Among the cell adhesion molecules, a great diversity of specific expression is observed (Figure 7). Members of the BEAT and SIDE family of adhesion molecules are expressed in a subtype-specific pattern with Beat-Ib enriched in dry cells and Beat-II enriched in hygrocool cells. Side is enriched in hot cells and Side-V in hygrocool cells of chamber I. Dscam2 and Dscam4 as well as the cadherin CadN2 are also expressed in unique patterns.

**Fig 7:**
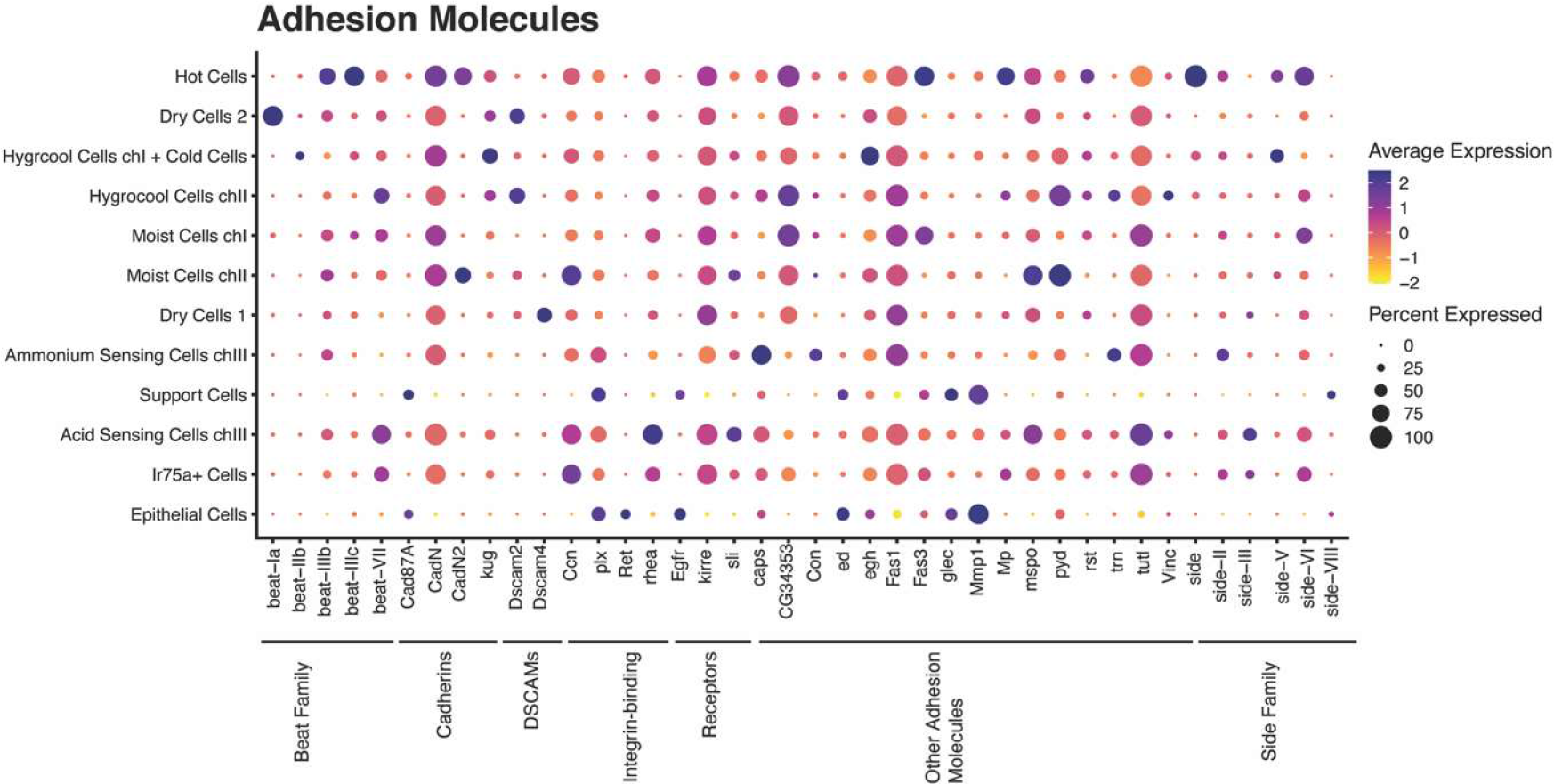
Cell Adhesion Molecules. Dot plot showing expression of selected cell adhesion molecules. Classification of genes is indicated by the black lines. The size of the dot indicates the fraction of cells in each cluster expressing the genes. The average gene expression for each cluster is shown as a heatmap.

### Enrichment of genes involved in cuticle development

The arista and hygrosensilla both have unique structural features of their overlying cuticle that may be related to its function. We therefore looked at genes involved in cuticle development in our dataset (Figure 8). Fourteen of the genes enriched are related to the GO term “cuticle development”. This includes transcription factors, secreted molecules, enzymes and receptors. Most cuticle development-genes are enriched in the support cells, which are the primary cells responsible for building and maintain the cuticle. However, a few cuticle-related proteins also show specific enrichment in HRNs. The cell adhesion molecule fred is specific for moist cells and the chitin deacetylase verm is specific for dry cells.

**Fig 8:**
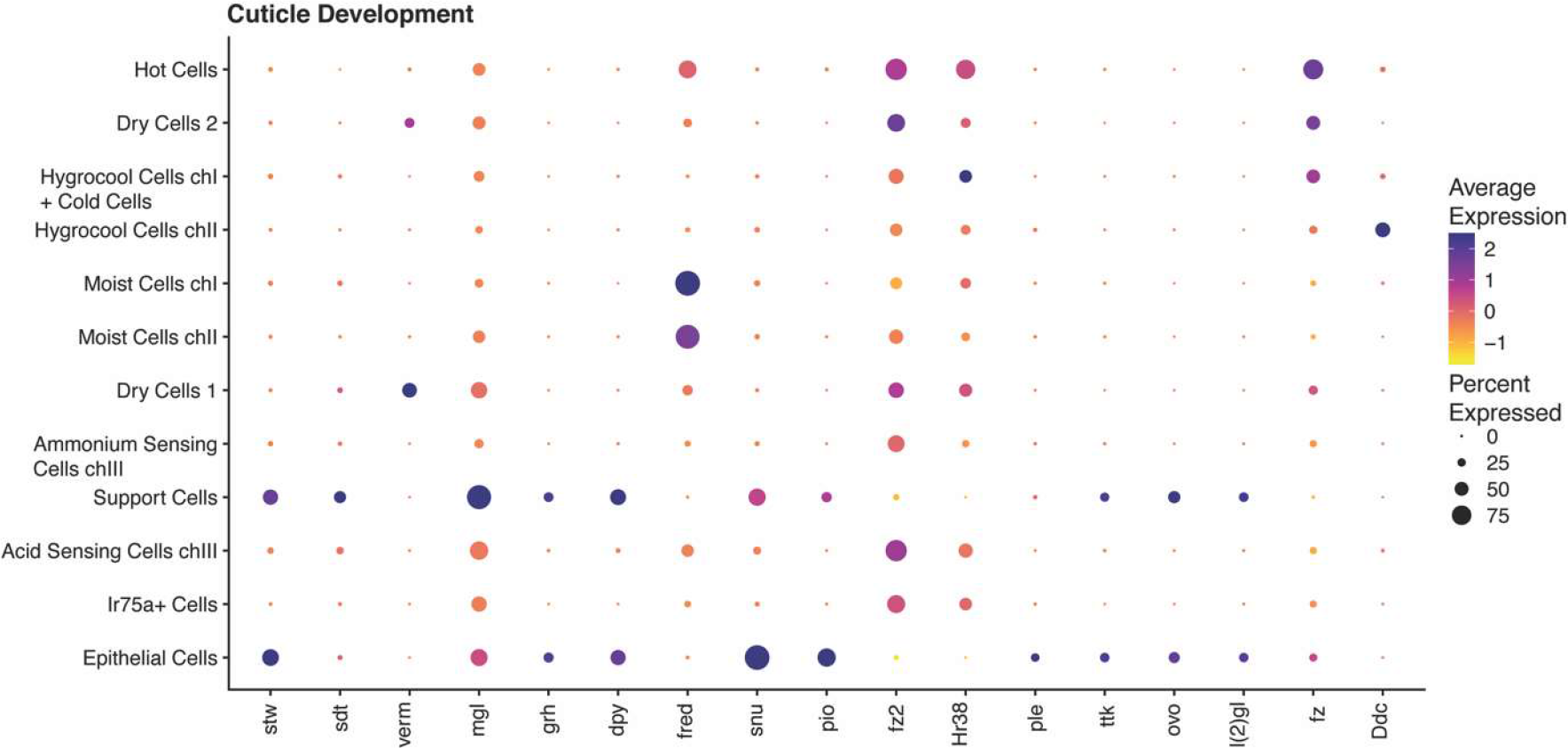
Cuticle Development. Dot plot showing expression of selected genes involved in cuticle development. Classification of genes is indicated by the black lines. The size of the dot indicates the fraction of cells in each cluster expressing the genes. The average gene expression for each cluster is shown as a heatmap.

## Discussion

The current study presents the first comprehensive transcriptomic atlas of the thermosensory and hygrosensory organ of adult *Drosophila melanogaster* with single-cell resolution. By performing unbiased clustering of more than 37,000 nuclei based on their patterns of gene expression, we identified 1411 nuclei that distributed into 12 distinct clusters of cells expressed in the sacculus and the arista. These clusters could be related to a known cell-type based on known markers and novel markers identified in this study. Furthermore, we identified and functionally annotated a large number of genes that were differentially expressed in each cluster compared to all other clusters. This dataset provides a resource for investigating genes involved in thermo- and hygrosensation in insects.

### Genetic profile of sacculus and arista cells

Our analysis revealed a notable genetic resemblance among adult TRNs, HRNs, and Ir75a-expressing cells in coeloconic sensilla. The neurons of the arista and sacculus are, together with neurons of coeloconic sensilla, specified from progenitor lineages expressing the transcription factor Atonal [32]. Among the neuron subtypes we have examined the cells responding to the same stimulus in arista, chamber I and II and among olfactory neurons are genetically the closest: the hot cells and cold cells of the arista, moist cells in chamber I and II, dry cells in chamber I and II, hygrocool cells in chamber I and II, and Ir75a/sacculus olfactory cells exhibit the highest degree of resemblance to one another.

Looking at the genes expressed specifically by these cell type we found that dry cells have enriched expression of nachralpha6, 5-HT7, Verm, Glucalpha, CG43795 and Oct-TyrR compared to other cells. In moist cells, we only find enriched expression of fred in both chambers. For the hygrocool cells clusters we cannot find any common unique genes between cells of chamber I and chamber II Of the genes enriched in arista and sacculus cells we find a large number of genes from the immunoglobulin superfamily. Dprs, DIPs, DSCAM, Beat and Side family members are unique markers for all populations of cells in the sacculus and arista. Members of these gene families interact with each other in a subtype-specific manner during development of the nervous system to establish neuronal connectivity [33–35]. The finding that this expression is maintained in the adult nervous system and suggests these genes have a role in the maintenance and structure of the nervous system [36].

Neuropeptides and neurohormones play key roles in the animal’s overall regulation of metabolism, physiology, and behavior, acting as global organizers at a higher hierarchical level [37]. We have found expression of several neuropeptide receptors in the sacculus and arista. It includes receptors involved in regulating circadian rhythms, feeding behavior and courtship such as Pdfr, sNPF-R and SPR [38–40]. These behaviors are known to be modulated by temperature and humidity. By identifying the subtype-specific expression of these receptors, the molecular and cellular components of temperature and humidity detection underlying the impact of temperature and humidity on these behaviors can be unraveled.

Other sensory processes like olfaction, gustation vision are dependent on genes involved in G-protein coupled receptors (GPCRs). As expected, genes involved in GPCR signaling are also differentially expressed in the hygrosensory and thermosensory neurons of the antenna. Nicotinic acetylcholine receptors (nAChRs) targets for antiparasitic drugs and insecticides [41,42]. In this dataset we find expression of nAChRalpha4 in all neuronal cells, nAChralpha6 as a distinct marker for the dry cell population and nAChRalpha7 in moist and hygrocool cells. Neuroligins are post-synaptic adhesion molecules that play a crucial role in synapse forming and maturation [43–45]. In *D.melanogaster* 4 neuroligins have been described: dnlg1-dnlg4 (further referred to as “Nlg1-4”) [46,47]. Previous studies describe severely impaired social and mating behavior in flies deficient for Nlg2 and Nlg4 [48,49]. Nlg4 is further associated with sleep regulation and flies deficient for Nlg4 show abnormal sleep cycle and circadian rhythm [49,50]. In *Apis mellifera* sensory deprivation in early development changes expression patterns of neuroligins [51]. Here, we find differential expression of Nlg2 (Dry Cells), Nlg3 (Moist Cells chII) and Nlg4 (Hygrocool chI+ cold cells and hot cells). To understand the role of neuroligins in humidity sensing and the underlying sensory processes further experiments are required. However, their presence in this group of sensory neurons is further evidence to their involvement in the plasticity of the sensory system.

### Concluding remarks

Overall, this dataset provides a valuable resource for future studies aiming to further understand the genetic basis of thermosensation and hygrosensation in insects. The identified cell clusters, marker genes, and differentially expressed genes offer new avenues for exploring the intricate mechanisms underlying sensory perception and behavior in response to temperature and humidity cues.

## Methods&Materials

### Animals

All used *Drosophila* strains were raised at 25 °C at a 12:12 h dark/light cycle on standard medium. The following strains were used (Bloomington id number in parenthesis): 5HT7-GAL4 (84592), nAChRalpha6-GAL4 (84665), Dh31-R-GAL4 (84625), Pdfr-GAL4 (84684), CG42458-GAL4 (67472), Ir68a-GAL4 (91305), Ir40a-GAL4 (41727), Octbeta2R-LexA (84437) and LexAOP-myr::smGdP-V5/UAS-myr::smGdP-HA (76358).

### Single nucleus RNA sequencing analysis

To analyze the transcriptomic profiles of the HRN and TRN population in the *Drosophila* antennae we used the 10x genomics antennal dataset from the recently published Fly Cell Atlas [15]. Quality control and subsequent analysis was done using the R package Seurat [16,17].

### Quality control

The antennal dataset was imported as a Seurat object and then processed using standard quality control metrics. The final data set retains cells expressing more than 200 but less than 2500 genes and less than 5% mitochondria.

This step ensures the quality of the used dataset and excludes possible doublets or damaged cells.

Next, the dataset was normalized by normalizing the gene expression measurement for each individual cell by the total expression, multiplied by a factor of 10,000 and the result log transformed. We then picked the 2000 most variable expressed genes for further analysis. The data was scaled and a principal component analysis (PCA) for the first 40 principal components was calculated.

### Nearest Neighbor Clustering and dimensionality reduction FCA full antennal data set

Unsupervised clustering of the data was done using the standard Seurat clustering approach. Briefly, this uses the Euclidian distance in the pre-defined PCA space (in this case 40 principal components) to compose a K-nearest neighbor (KNN) graph with borders enclosing communities of cells with similar expression patterns. To optimize the standard modularity functions the Louvain algorithm was applied using a resolution of 0.5.

To visualize the resulting clusters the non-linear dimensional reduction UMAP was used with the previously selected 40 PCs.

### Assigning cluster identities and identifying the HRNs

To identify the clusters containing the HRN populations we initially searched for known marker genes in the whole antennal neuronal dataset. We then confirmed these findings by calculating the most differentially expressed genes in those clusters, only reporting the positive markers. Based on this we selected clusters of interest for further analysis.

In the next step we extracted those clusters from the whole data set and ran all commands from normalization to clustering again on this subset of cells. Here, we again used 40 principal components and a resolution of 0.5, resulting in 12 clusters.

To assign an identity of all resulting clusters we searched for expression of known marker genes for the expected cell types, allowing us to classify all clusters.

### Separating the hygrocool cells and temperature cells

In the sub-clustering containing the sacculus and arista cells we can find 3 clusters with assumed thermosensitive identity: arista hot cells, hygrocool cells chII and hygrocool cells ch I + arista cold cells.

To separate the hygrocool cells from the arista cold cells, we extracted those three clusters and followed the Seurat pipeline from normalization to NN clustering. Using 10 principal components and a resolution of 0.7 results in 4 clusters that can be identified as hygrocool cells chI, hygrocool cells chII, hot cells and cold cells.

### Selection and classification of differentially expressed features

Each cluster is defined by a set of differentially expressed genes (or biomarkers). Using the Seurat ‘FindMarkers’ function (with min.pct set to 0.25, reporting only positive markers) returns the markers for each cluster, compared to all remaining cells.

To find the top markers, they were sorted by the average log2FC and the top5, top50 and top100 genes were selected, respectively. The genes listed as top 100 markers were classified with GLAD, Flybase gene group and Gene Ontology term using the PANGEA tool [31].

### Visualization of gene expression

To visualize the expression of marker genes across the different clusters, we are using dot plots. In this style of plot, the average expression of a gene in the cluster is using a heatmap of colors. The size of the dot indicates the percentage of cell express the gene within the cluster. To visualize our data we used the scCustomize package for R [52].

### Immunoflourescent stainings

Flies were anesthetized and the antenna dissected. The antennae were fixed in 2% paraformaldehyde (PFA, Electron microscopy studies) in phosphate-buffered saline with 0,5% Triton X-100 (PBST, Sigma-Aldrich) for 55min at RT and subsequently washed with PBST for 4 x 10min at RT on a shaker.

Blocking was done using 5% goat serum (Thermofisher) in PBST for 1.5h at RT and primary antibody was added (GFP A11122 or A11070 Invitrogen, HA-Tag C29F4 Cell signaling technology) diluted 1:300 in 5% goat serum in PBST. The primary antibody is incubated for 4h on a shaker at RT for 4h and afterwards for 48h on 4°C.

After incubation the samples are washed in PBST 5×15min on a RT shaker secondary antibody (Alexa Fluor 488 goat anti-rabbit IgG A11008 Invitrogen, V5-TAG:DyLight550 MCA1360D550GA Biorad,) diluted 1:500 in 5% goat serum in PBST is added to the samples. The secondary antibody is incubated for 4h at RT and afterwards for 48-72h on 4°C. Samples are washed in PBST 5×15min on a shaker at RT and afterwards mounted with RapiClear (SunJin Lab, Hsinchu, Taiwan, RC149002). Imaging was performed on a Leica SP8 confocal microscope with a 63x, 1.4 NA objective at a resolution of 512×512.

## ACKNOWLEDGMENTS

We wish to thank the NBIS for their support on this project. The authors thank Marcus Stensmyr for comments on discussion of this project. This project was funded by the WennerGren foundation, Formas – a Swedish Research Council for Sustainable Development, the Swedish Research Council, the Crafoord foundation and the Jaenssons foundation. AC and JR are financially supported by the Knut and Alice Wallenberg Foundation as part of the National Bioinformatics Infrastructure Sweden at SciLifeLab.

## Author Contribution

AE designed the study. KC performed the transcriptomic analysis with input from AC and JR. VA performed the immunostainings. AE and KC wrote the manuscript with input from all co-authors.

## Data availability

Data was from https://flycellatlas.org/. In this study we used the 10x antennal data set in loom-file format. Raw data is available

