## Supplemental figures for "Genetic atlas of hygro- and thermosensory cells in the vinegar fly *Drosophila* melanogaster"

### Supplementary Figures

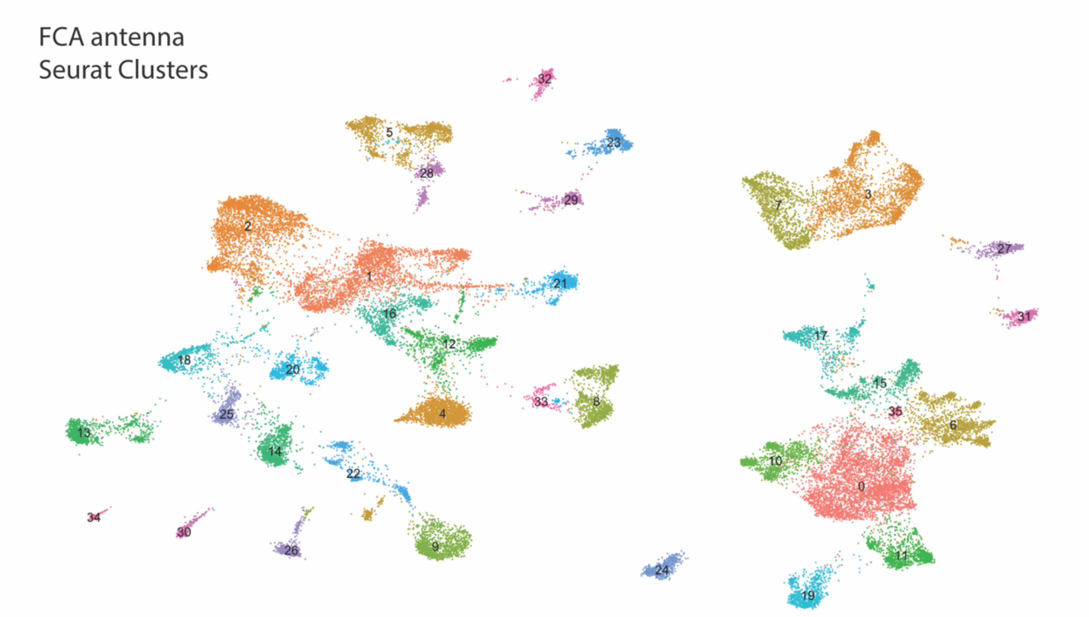


0: ORN (Orco+, Or35a+, Or43b+)
1: epthelial cells (snu+, osy+, grh+)
2: epithelial cells (snu+, osy+, grh+)
3: Johnston's organ (nan+, lav+, Dnah3+, Dnai2+)
4: ORN (Orco+)
5: epithelial cells (snu+, osy+)
6: ONR (Ir84+, Ir31a+, Ir76a/b+, Or35a+)
7: Johnston's organ (nan+, lav+, Dnah3+, Dnai2+)
8: epithelial cells (shg+)
9: epithelial cells (snu+, osy+)
10: ORN (Orco+, Or42b+)
11: ORN (Orco+, Ir47b+)
12: epithelial cells (snu+, osy+)
13: epithelial cells (snu+, osy+, sv+)
14: epithelial cells (sv+)
15: sacculus (Ir40a+, Ir64a+, Ir21a+), arista (Gr28b+, Ir21a+), acid sensing (Ir64a+, Ir75a/b/c+)
16: epithelial cells (snu+, osy+, grh+)
17: unannotated neurons (nSyb+)
18: epithelial cells (snu+, osy+, grh+)
19: ORN (Or67d+)
20: epithelial cells (snu+, osy+, sv+)
21: glia cells (repo+)
22: epithelial cells (snu+, osy+, sv+)
23: glia cells (repo+), hemo-cytes (hml+, NimC1+)
24: ORN (Ir75d+)
25: epithelial cells (sv+)
26: epithelial cells (sv+)
27: ORN (Gr21a+, Gr63a+)
28: epithelial cells (snu+, osy+)
29: glia cells (repo+)
30: glia cells (repo+)
31: ORN (Orco+, Or65a+)
32: muscle cells (Mhc+)
33: epithelial cells (snu+, osy+, sv
34: epithelial cells (shg+)
35: ammonium sensing cells (Rh50+, Amt+)

SuppFig1: Annotation of full antennal dataset of the FlyCellAtlas. Clustering of the FCA antennal dataset using the Seurat pipeline (see Material&Methods for details) resulted in 36 clusters. Based on top marker gene expression we assigned identities to each cluster. These identities are corresponding to the clusters found in the original publication. The list shows each cluster with assumed identity and the gene on which identification is based. All neuronal clusters are also positive for the neuronal marker nSyb, even if not specified. Cluster 17 remains unannotated, based on expression of the neuronal marker nSyb it is assumed to have neuronal identity.

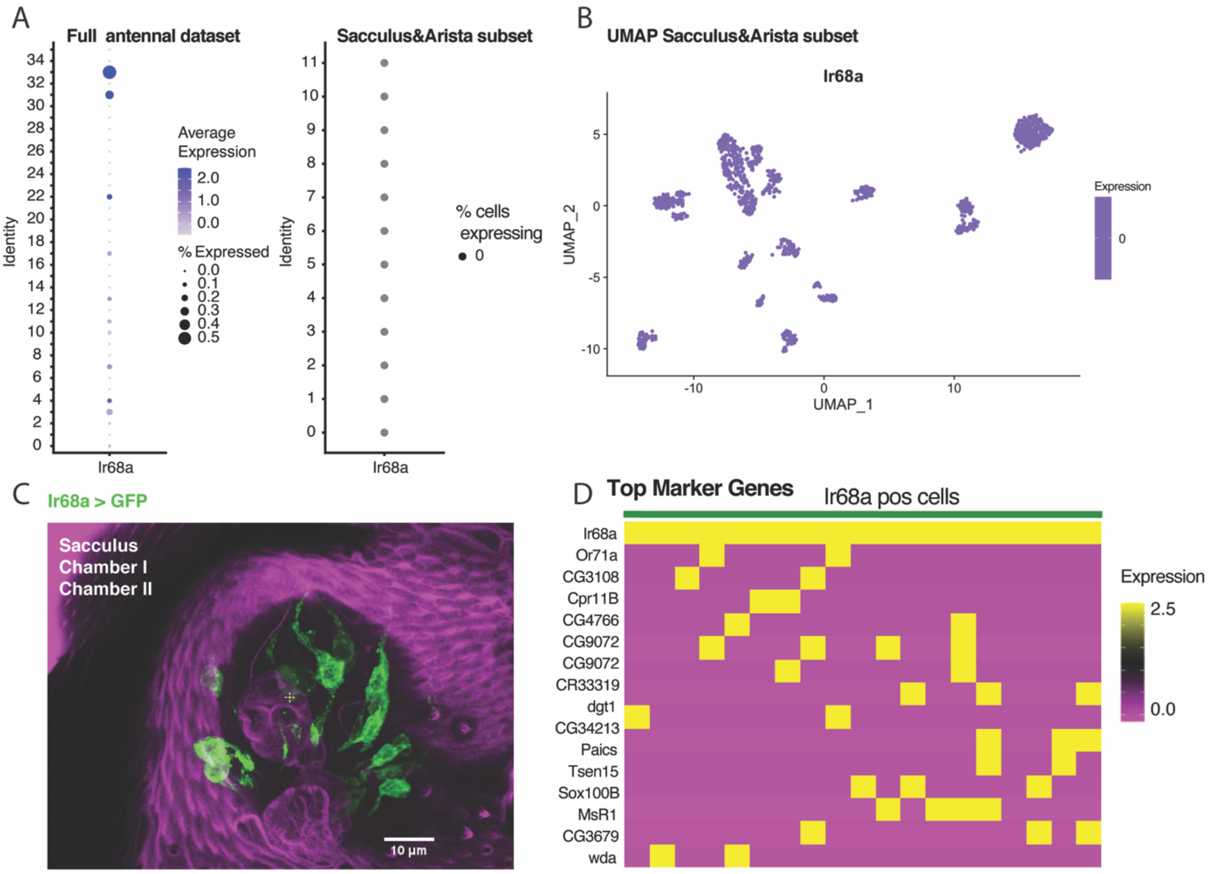


SuppFig2: Expression of Ir68a. A) Expression of Ir68a in the FCA full antennal dataset and the sacculus and arista cells. No clusters show a high percentage of cells expressing Ir68a. No expression of Ir68a can be found in the arista and sacculus cells. B) UMAP of Ir68a expression in sacculus and arista cells. No expression of Ir68a can be found. C) Immunostaining of Ir68a. An Ir68a-Gal4 line was crossed to a GPF reporter line (see Material&Methods for more details on fly strains). Subsequent GPF staining shows clear expression of Ir68a in both chamber I and II of the sacculus. D) Marker genes for all cells expressing Ir68a.  Selecting all cells expressing Ir68a from the FCA full antennal dataset results in a group of 19 cells. Results of finding marker genes of this group opposed to all other cells in the FCA are shown here. Based on this marker expression we assume that the 19 cells expressing Ir68a are not a homogenous group.

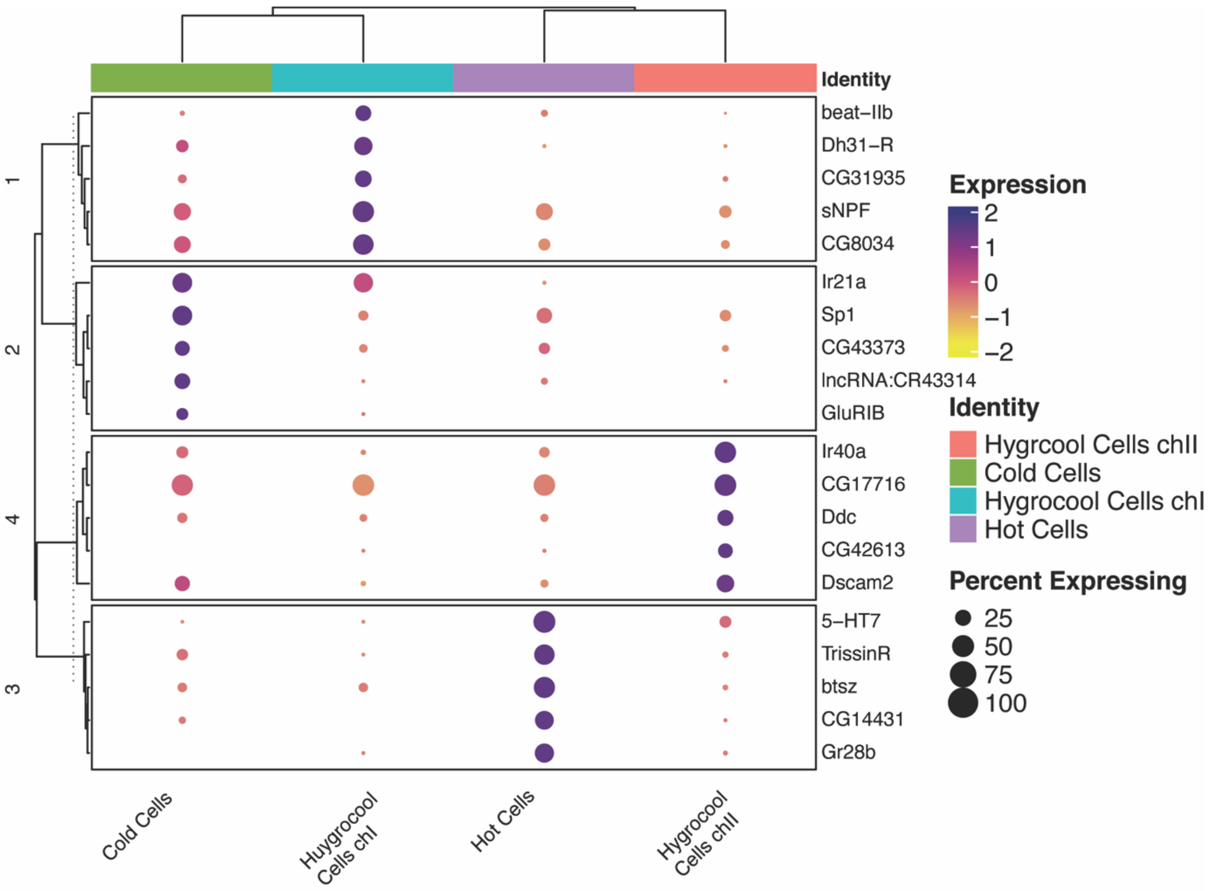


SuppFig3: Sub-clustering the temperature sensitive cells from cluster 8 (hygrocool cells chII), 9 (hygrocool cells chI + cold cells) and 11 (hot cells). Clustered dot plot shows the expression of top marker genes for each cluster. The dendrogram indicates the relationship between clusters based on this expression. Sub-clustering results in 4 clusters: hot cells, cold cells, hygrocool cells chI and hygrocool cells chII.

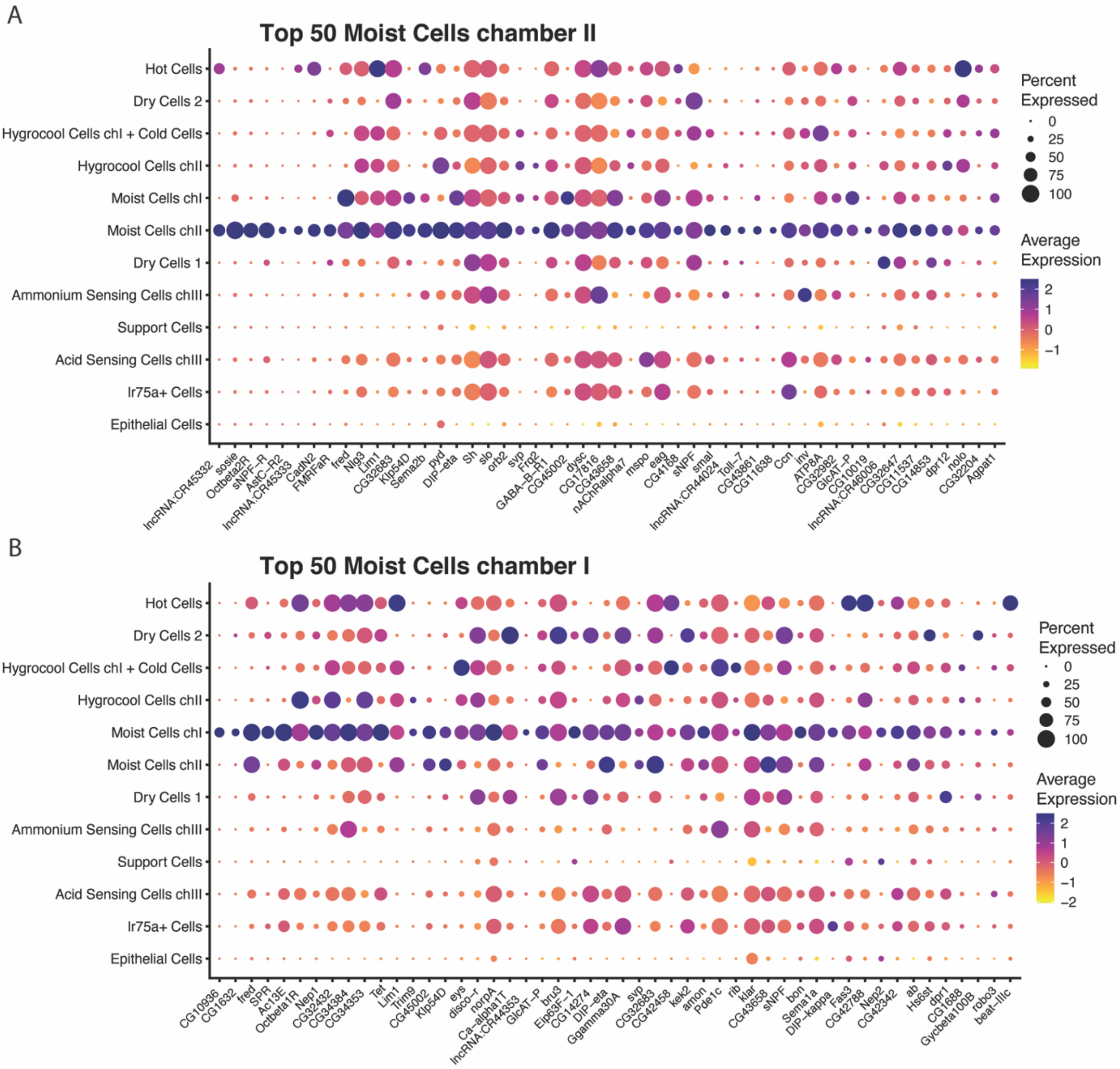


SuppFig4: Top 50 Markers for moist cell clusters. Dot plot showing expression of top 50 markers for each cluster. Classification of genes is indicated by the black lines. The size of the dot indicates the fraction of cells in each cluster expressing the genes. The average gene expression for each cluster is shown as a heatmap. A) Top 50 markers for moist cells chII. B) Top 50 markers for moist cells ch I.

A

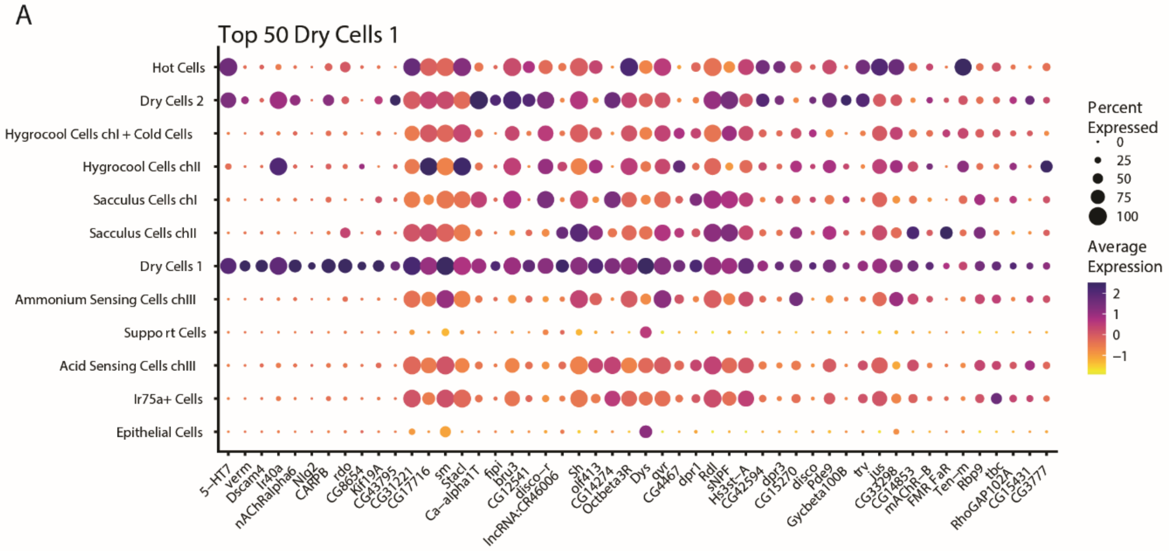


B

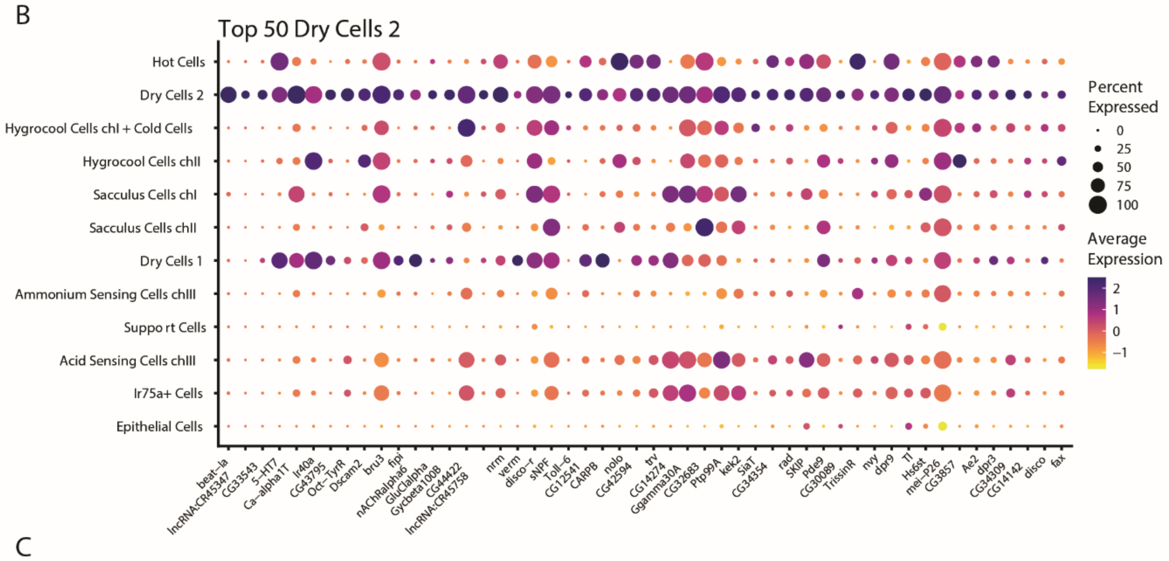


C

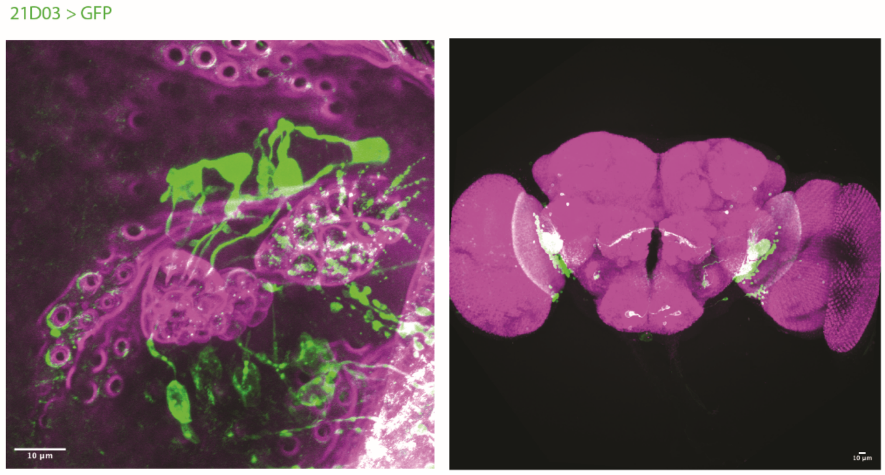


SuppFig5: Top 50 Markers for dry cell cluster. Dot plot showing expression of top 50 markers for each cluster. Classification of genes is indicated by the black lines. The size of the dot indicates the fraction of cells in each cluster expressing the genes. The average gene expression for each cluster is shown as a heatmap. A) Top 50 markers for dry cell cluster 1.

B) Top 50 markers for dry cell cluster 2. C) Immunostaining of GMR21D03>CD8::GFP antenna and brain. Brain images from Flylight project and used under CC BY 4.0-licence [53].

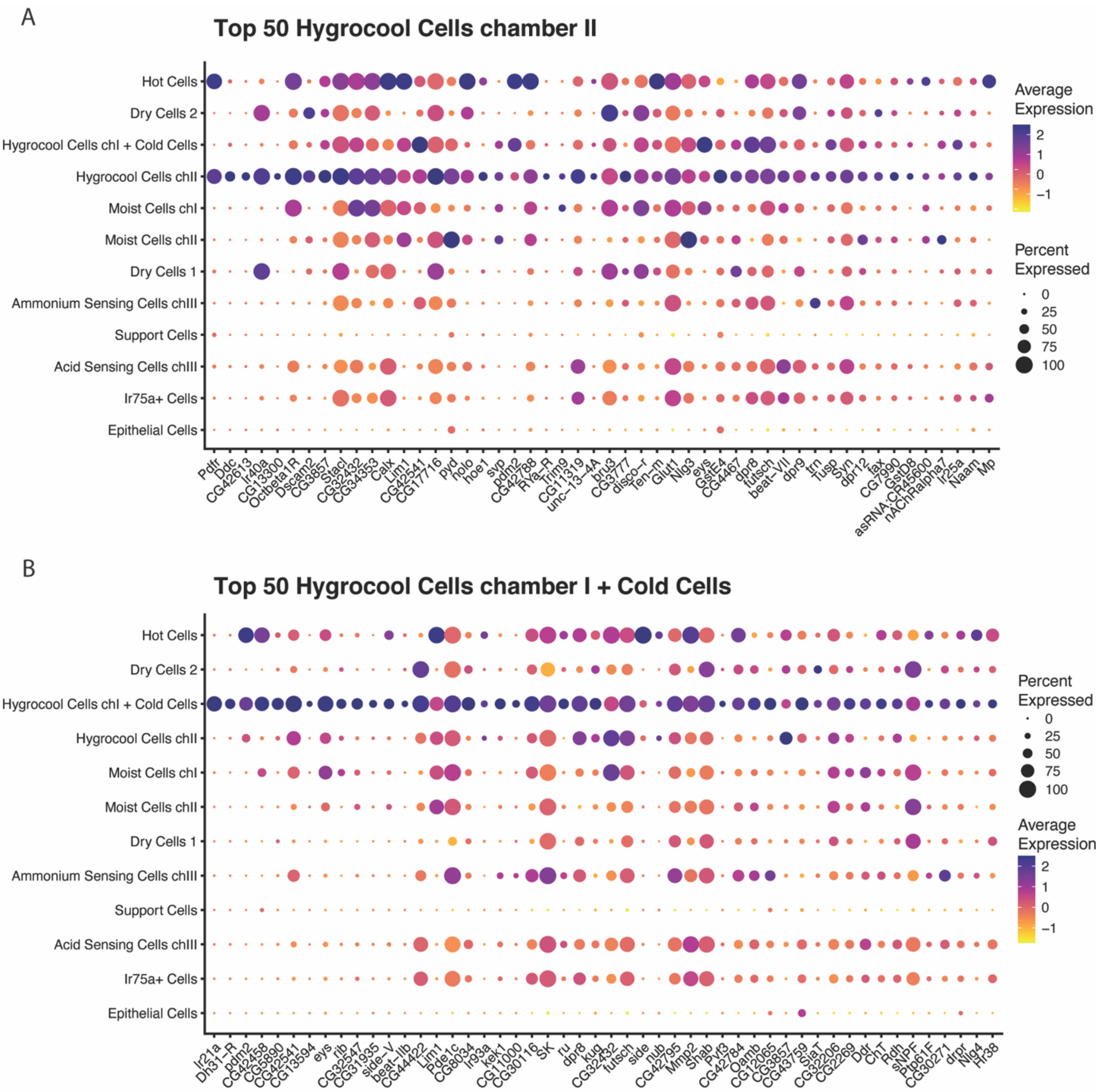


SuppFig6: Top 50 Markers for hygrocool cell cluster. Dot plot showing expression of top 50 markers for each cluster. Classification of genes is indicated by the black lines. The size of the dot indicates the fraction of cells in each cluster expressing the genes. The average gene expression for each cluster is shown as a heatmap. A) Top 50 markers for hygrocool cells chII. B) Top 50 markers for hygrocool cells ch I + cold cells.

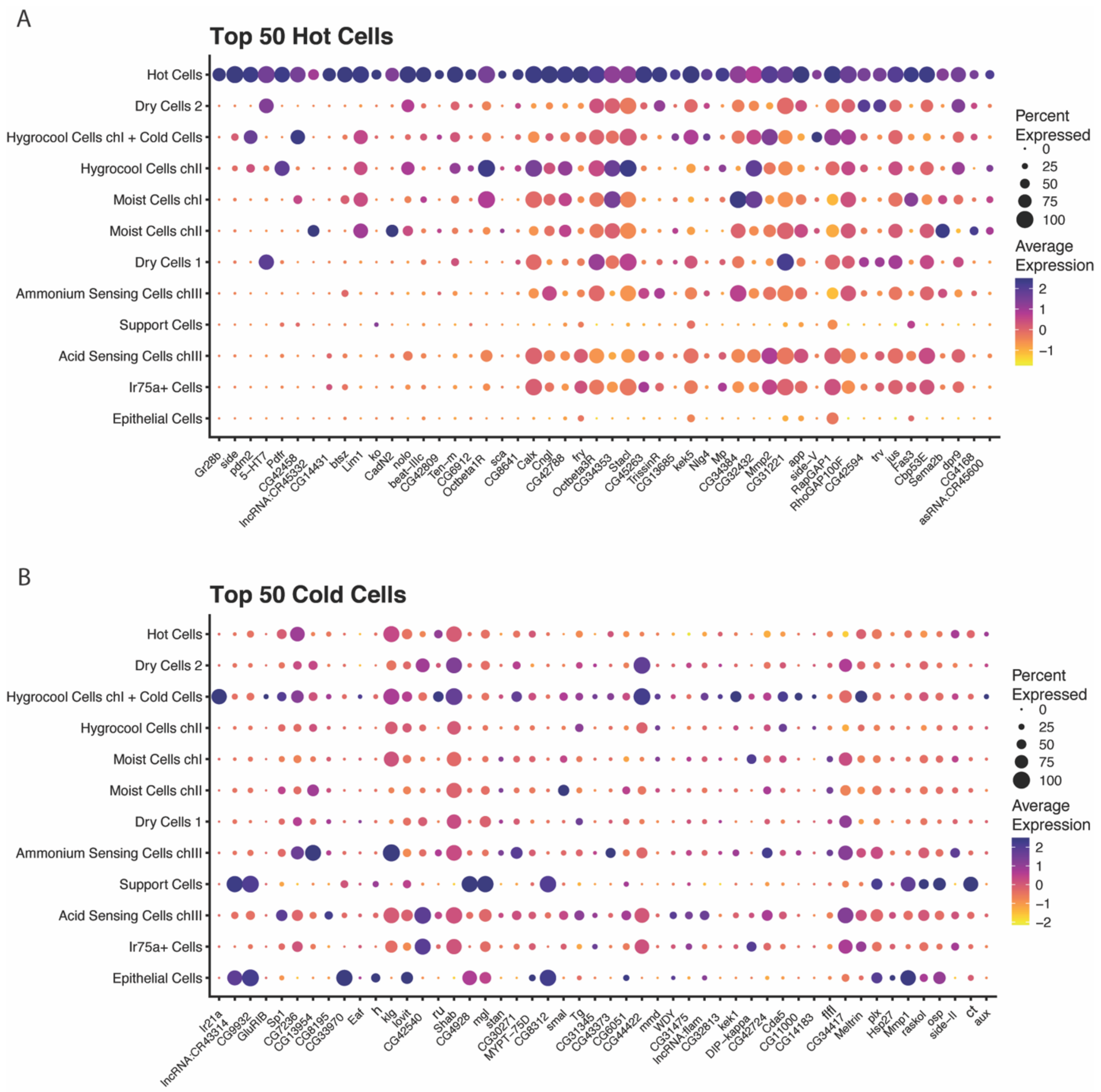


SuppFig7: Top 50 Markers for arista temperature cells. Dot plot showing expression of top 50 markers for each cluster. Classification of genes is indicated by the black lines. The size of the dot indicates the fraction of cells in each cluster expressing the genes. The average gene expression for each cluster is shown as a heatmap. A) Top 50 markers for hot cells. B) Top 50 markers for cold cells.

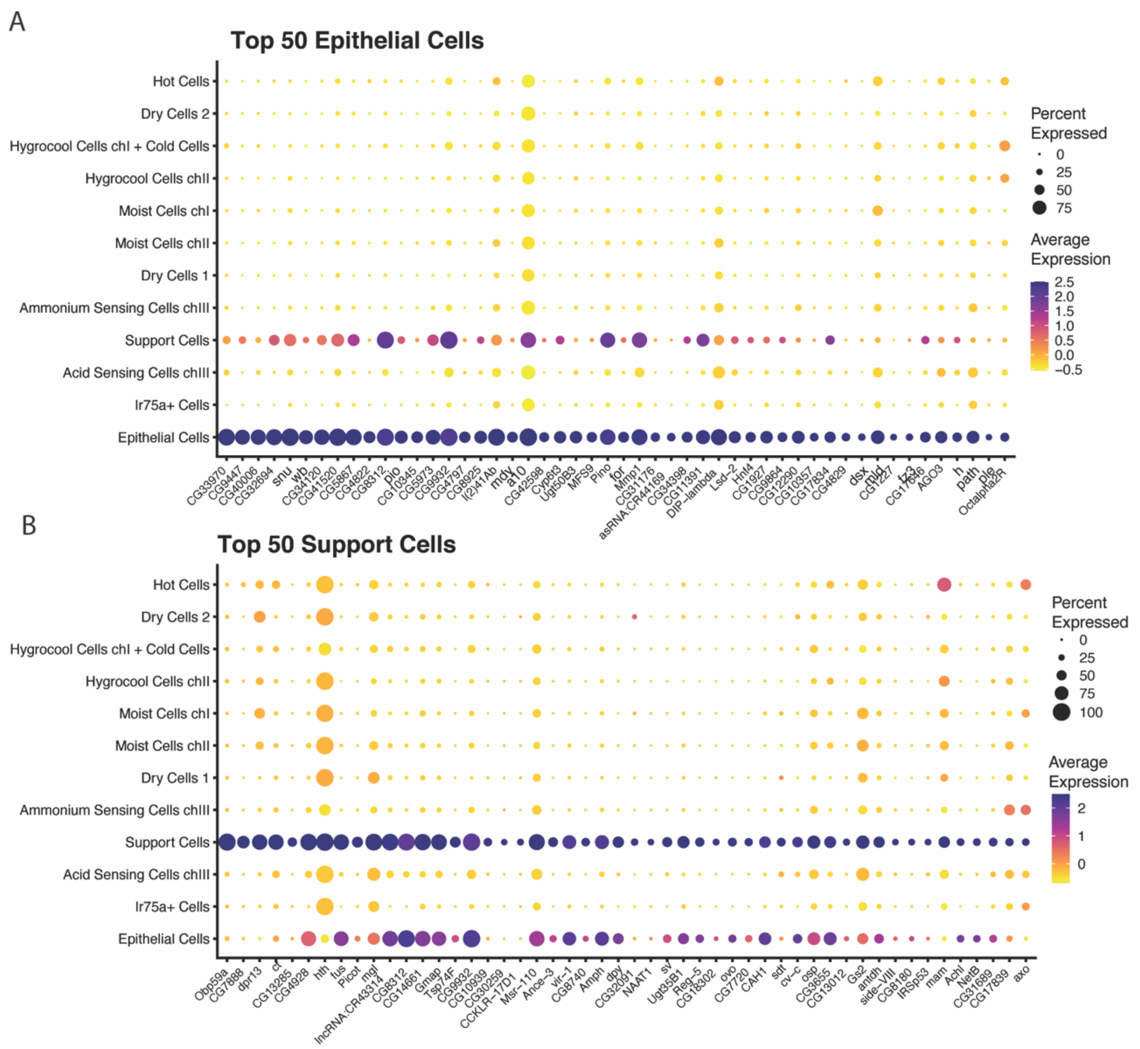


SuppFig8: Top 50 Markers for support cells and epithelial cells. Dot plot showing expression of top 50 markers for each cluster. Classification of genes is indicated by the black lines. The size of the dot indicates the fraction of cells in each cluster expressing the genes. The average gene expression for each cluster is shown as a heatmap. A) Top 50 markers for epithelial cells. B) Top 50 markers for support cells.
